# Development of a Low-Cost System for Simultaneous Longitudinal Biological Imaging

**DOI:** 10.1101/2021.05.17.443454

**Authors:** Victoria T. Ly, Pierre V. Baudin, Pattawong Pansodtee, Erik A. Jung, Kateryna Voitiuk, Yohei M. Rosen, Helen Rankin Willsey, Gary L. Mantalas, Spencer T. Seiler, John A. Selberg, Sergio A. Cordero, Jayden M. Ross, Marco Rolandi, Alex A. Pollen, Tomasz J. Nowakowski, David Haussler, Mohammed A. Mostajo-Radji, Sofie R. Salama, Mircea Teodorescu

## Abstract

Simultaneous longitudinal imaging across multiple conditions and replicates has been crucial for scientific studies aiming to understand biological processes and disease. Yet, imaging systems capable of accomplishing these tasks are economically unattainable for most academic and teaching laboratories around the world. Here we propose the Picroscope, which is the first low cost system for simultaneous longitudinal biological imaging made primarily using off-the-shelf and 3D-printed materials. The Picroscope is compatible with standard 24-well cell culture plates and captures 3D z-stack image data. The Picroscope can be controlled remotely, allowing for automatic imaging with minimal intervention from the investigator. Here we use this system in a range of applications. We gathered longitudinal whole organism image data for frogs, zebrafish and planaria worms. We also gathered image data inside an incubator to observe 2D monolayers and 3D mammalian tissue culture models. Using this tool, we can measure the behavior of entire organisms or individual cells over long time periods.

## INTRODUCTION

Monitoring and handling live tissues and cell cultures as well as analyzing their secreted contents are essential tasks in experimental biology and biomedicine. Advances in microscopy have revolutionized biological studies, allowing scientists to perform observations of cellular processes and organisms’ development and behaviors. Imaging has been pivotal to uncovering cellular mechanisms behind biological processes^1^. Several options exist on the market to perform longitudinal imaging of biological materials. These range from super-resolution microscopes that allow the imaging of individual biomolecules^2, 3^ to conventional benchtop microscopes, which are common in academic research^3–7^, industrial^8, 9^ and teaching laboratories^10^. When deciding between the different technologies for longitudinal live tissue imaging, several factors need to be considered in the experimental design. The image acquisition speed of the microscope should be sufficient for the phenomenon being studied. The microscope should be able to acquire images without damaging or disturbing the specimen, such as photobleaching. The microscope should be capable of imaging in the environmental conditions needed for the desired experiment, including temperature, light, and humidity. The resolution of the microscope should be sufficient to view the phenomenon being studied. When scaling to simultaneous mutli-well longitudinal tissue imaging it is also important that the apparatus not be bulky or expensive. It has been challenging to meet all of these criteria.^11^.

The use of open-source technology, including 3D printers, laser cutters, and low-cost computer hardware, has democratized access to rapid prototyping tools and dramatically increased the repertoire of biomedical equipment available to laboratories around the world^12, 13^. Through rapid prototyping and the use of open source platforms, the technology can be replicated and quickly improved^14, 15^. 3D printer technology has been applied to several fields in biomedicine, including biotechnology^16^, bioengineering^17, 18^, and medical applications including fabrication of tissues and organs, casts, implants, and prostheses^19^. Existing 3D printed microscopes range in complexity from simple low cost systems with pre-loaded imaging modules^20^ to portable confocal microscopes capable of imaging individual molecules^21^ and even 3D printed microfluidic bioreactors^22^.

The majority of low-cost 3D printed microscopes are not intended for longitudinal imaging of simultaneous biological cultures (e.g., multi-well, multi-week biological experiments). They usually have a single imaging unit^5, 17, 23–32^ or perform confocal,^21^ and even light-sheet imaging^25^. Other systems have taken advantage of one camera attached to a gantry system to perform imaging of multiple experimental replicates^33–35^. Few 3D-printed microscopes have been developed that perform multi-well imaging with medium throughput^34, 36^. Several biological applications exist that would greatly benefit from multi-well, multi-week simultaneous imaging, as it allows for concurrent interrogation of different experimental conditions and the inclusion of biological replicates. These include cell culture applications, in which 2D and 3D culture models can be tracked over multi-week periods, as well as developmental and behavioral biology experiments in which multi-week tracking could be performed on whole organisms.

Here, we report a new simultaneous multi-well imaging system (the Picroscope), which features a low cost per well ($83) and performs longitudinal brightfield z-stack imaging of 24 well cell culture plates. Images are uploaded to a server as they are captured allowing the users to view the results in near real time. We used this system to longitudinally track different animal models of development and regeneration, including *Xenopus tropicalis* (frogs), *Danio rerio* (zebrafish), and planaria worms. Finally, we demonstrate this system’s versatility by imaging human embryonic stem cells and 3D cortical organoids inside a standard tissue culture incubator. We demonstrate that the Picroscope is a robust low-cost, versatile multi-well imaging system for longitudinal live imaging biological studies.

## RESULTS

### System design

Typically 3D manufacturing has two main approaches: additive and subtractive, where both require dedicated equipment. In the past some of these devices were limited to specialized manufacturing facilities. Over the past couple of decades, 3D manufacturing went thorough a revolution. Equipment such as 3D printers and computer numerical control (CNC) machinery has become affordable and ubiquitous in most university engineering laboratories. Research in the areas of labs-on-chip, optofluidics, microscopy, in combination with developments in consumer oriented tools for “makers”, has the potential to democratize access to cell biology based research.^17, 28^. Laboratories are now able to more easily develop custom devices which can be shared with the greater research community as open source projects^17^ (Table 1). Laboratories are able to develop custom devices which are often shared with the greater research community as open source projects^17^ (Table 1). The Picroscope is one such device.

**Table 1.** Comparison between 3D printed microscopes

| Low Cost Maker Microscope | Single or Multi-Camera | Price per Camera |
| --- | --- | --- |
| Foldscope<sup>30</sup> | Single | \$1 |
| LudusScope<sup>31</sup> | Single | \$30-100 |
| FlyPi<sup>23</sup> | Single | \$120.93 |
| A versatile and customizable low-cost 3D-printed open standard for microscopic imaging<sup>25</sup> | Single | \$120.92 - \$483.69 |
| Incu-Stream 1.0<sup>35</sup> | Single | \$184 |
| Portable, Battery-Operated, Low-Cost, Bright Field and Fluorescence Microscope<sup>3</sup> | Single | \$240 |
| Microscopi<sup>7</sup> | Single | \$419.06 |
| The Incubot<sup>34</sup> | Single | \$1,208.95 |
| An inexpensive system for imaging the contents of multi-well plates<sup>33</sup> | Single | \$3400 |
| Single-molecule detection on a portable 3D-printed microscope<sup>21</sup> | Single | \$N/A |
| A mini-microscope for in situ monitoring of cells<sup>24</sup> | Single | \$N/A |
| Low-cost, sub-micron resolution, wide-field computational microscopy using opensource hardware<sup>32</sup> | Single | \$N/A |
| EmSight<sup>36</sup> | Single | \$N/A |
| ***The Picroscope*** | ***Multi-Camera*** | ***\$83 (\$2000 for 24 cameras)*** |

The Picroscope is a programmable, data rich, sensor-per-well simultaneous imaging system for longitudinal brightfield imaging to automate microscopy (Figure 1). The system simultaneously images in each one of the 24 wells multiple focal planes (the resolution of the “z-stack” can be remotely changed) several times every hour for weeks, a frequency impractical to perform manually. The instrument is made from off-the-shelf components (e.g., lenses, motors, cameras, Arduino^37^, Raspberry Pi^38^, MakerBeam aluminum extrusions^39^), and 3D printed Polylactic acid (PLA) components with 100% infill (the percentage of the internal part of the piece that is occupied by the printing material).

**Figure 1.**
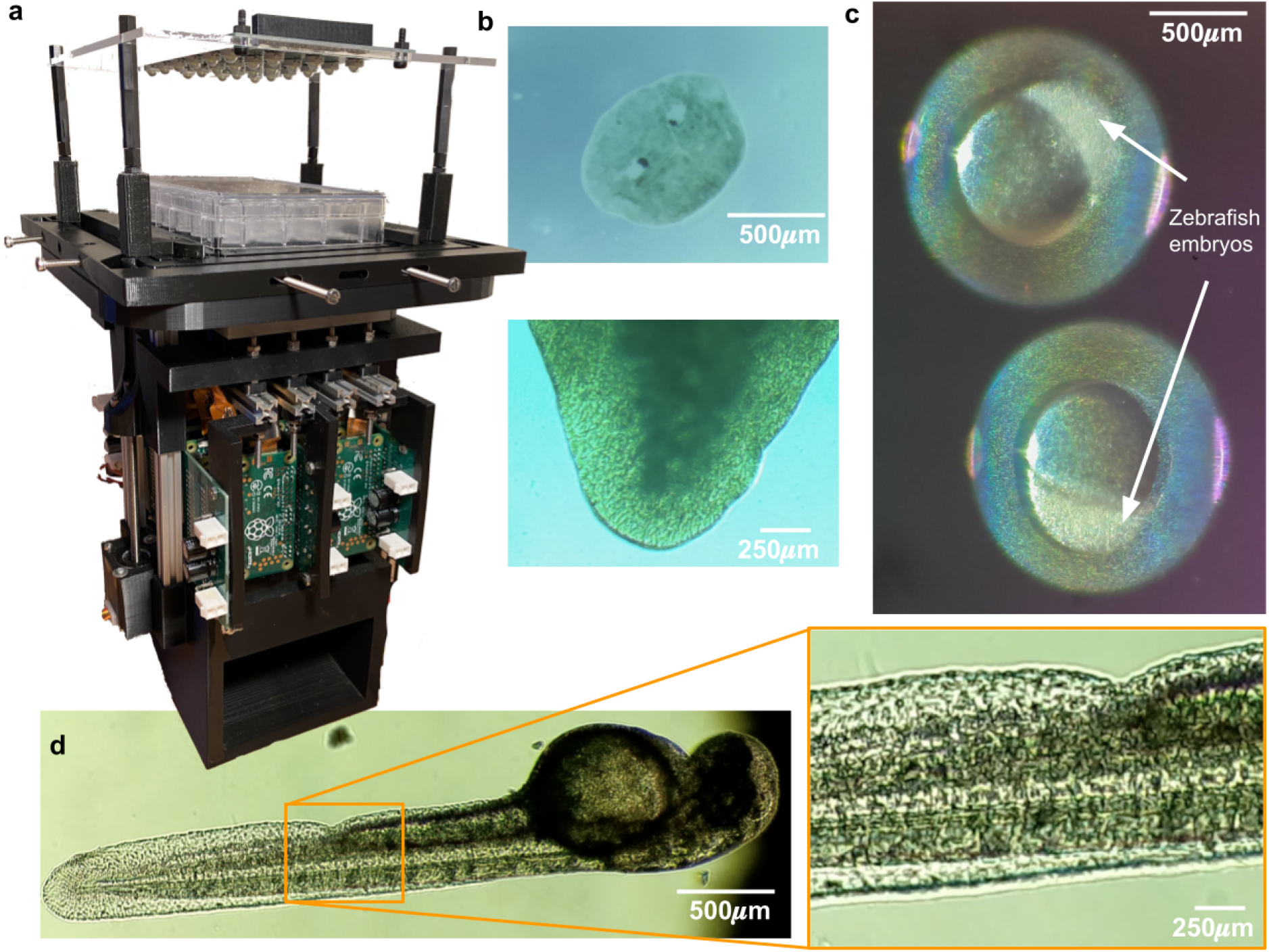
Development of a low-cost system for simultaneous longitudinal biological imaging. **a** The Picroscope fits a standard 24 well plate, it is controlled remotely and images can be accessed through a web browser. **b-d** Applications of the Picroscope to longitudinal imaging of developmental biology and regeneration. **b** Regeneration of planaria worms Dugesia tigrina. **c** Zebrafish embryonic development at oblong stage. **d**. Zebrafish embryo at 48 hours post fertilization. Video S1.

The picroscope is designed to illuminate the samples using one or multiple lighting sources (from above or below a standard 24 well cell culture plate)(Figure 2). Diffused illumination from below results in images that show contours and surface features. Illumination from above typically results in more visible detail and can show internal structures if the sample is sufficiently translucent. The flexibility of using different illumination techniques emulates commercial brightfield microscopes. The 3D printed plate holder (2 in Figure 2) supports the biological sample during an experiment. For easy alignment the holder is attached to a xy sliding stage that consists of two interconnected linear stages (Figure 2f). The inner stage translating along the y axis uses 8 leaf springs to connect a central piece holding the 24-well plate with 4 rigid elements surrounding it. The outer stage translating along the x axis uses 8 additional leaf springs to connect the inner stage with the outside 4 rigid elements, two of them being connected to the picroscope frame using 4 screws (18 in Figure 2). While each stage is flexible along one axis (x or y), together they can slide along both, x and y axes. Each stage is actuated by two adjustment screws depicted as gray arrows in Figure 2f.

**Figure 2.**
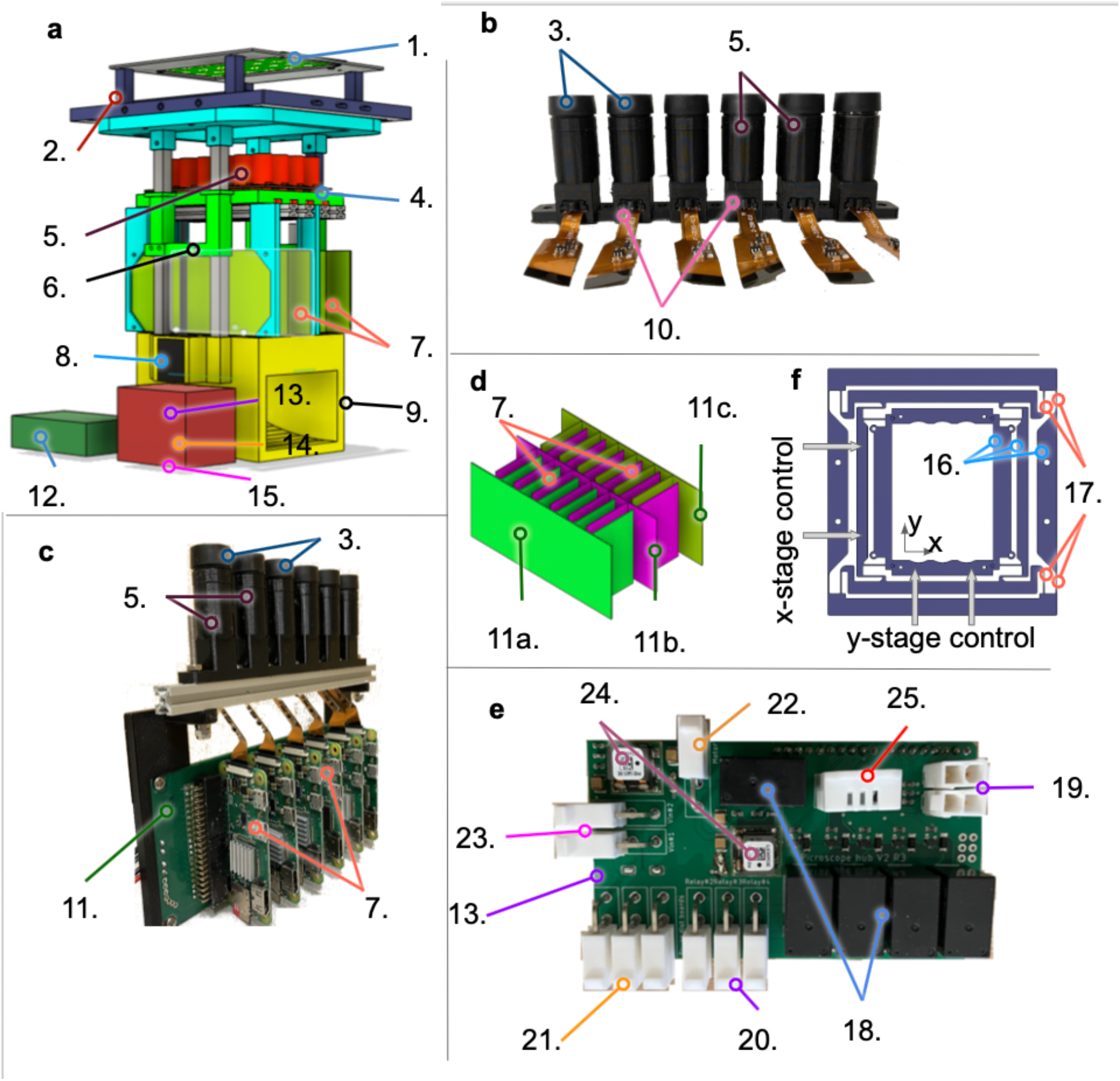
The Picroscope. **a** Physical representation of the proposed imaging system. **b** one line of independent cameras. **c** An integrated rack of cameras and raspberry pi board computers. **d** The interlacing strategy of four independent racks of power distribution boards. **e** The Raspberry pi Hub and Arduino Uno, Motor driver and custom relay board **f**The XY adjustment stage. 1 = Over-the-plate illumination board, 2 = 3D printed Cell Culture Plate Holder & XY stage, 3 = Lenses, 4 = Illumination Board from below, 5 = 3D Printed Camera Bodies, 6 = 3D Printed Elevator, 7 = Raspberry Pi 0Ws, 8 = Motors, 9 = Base, 10 = Raspberry Spy Cameras, 11 = Interface Board a. row 1, b. rows 2 and 3 c. row 4, 12 = Pi Hub – Raspberry Pi 4, 13 = Custom Relay Board, 14 = Adafruit Motor/Stepper/Servo Shield for Arduino v2, 15 = Arduino Uno, 16 = Leaf Springs, 17 = Rigid Elements, 18 = Relays, 19 = Limit switches connectors, 20 = Power distribution board connectors, 21 = Light board connectors, 22 = Motor power connector, 23 = 12 V power source, 24 = Voltage regulators, 25 = Temperature & Humidity sensor

The imaging unit comprises 24 independent objectives attached to a vertical sliding stage (“elevator piece” in 2a) using 4 makerbeam vertical columns and 2 Nema-11 stepper motors (3.175*µ*m Travel/Step). The fine threads are necessary for focusing of specific biological features and collecting z-stack imaging (Figure3). The objectives are distributed on 4-rows and 6-columns to match a standard 24 well culture plate. Each objective consists of a 3D printed camera body that hosts a 5 MegaPixel (5MP) camera (Spy Camera for Raspberry Pi0W, with a 1.4 µm × 1.4 µm pixel pitch)^40^ and an off-the-shelf Arducam 1/2” M12 Mount 16 mm Focal Length. Each objective is controlled by a single board computer (Raspberry Pi 0Ws), which is connected to an individual slot on one of the 3 custom-made power distribution boards (2c and D). All 24 single board computers (Raspberry Pi0W) computers communicate to a hub board computer (Raspberry Pi 4) that manages the images and autonomously uploads them to a remote server. The hub single board computer has the MIPI CSI-2 camera port and is connected to an Arduino Uno, which has a motor shield attachment, to controls the motors and lift the elevator piece (2e). As a safety feature, the system also includes a custom-made Relay Board that is attached to the Arduino and motor driver stack. The relay board provides control of the LED boards and in the event of an overheat allows us to shut down the system, protecting the system and the biological sample. After each set of pictures, the imaging unit returns to the lowest (“park”) position, which is determined by a limiting switch attached to the elevator unit. The entire system sits on a 3D printed base, that includes a fan for heat dissipation. During the course of an experiment, the pictures are autonomously uploaded on a remote computer/server using the ethernet connection of the hub computer board, where they can be viewed or processed in near real time (see figure 4).

**Figure 3.**
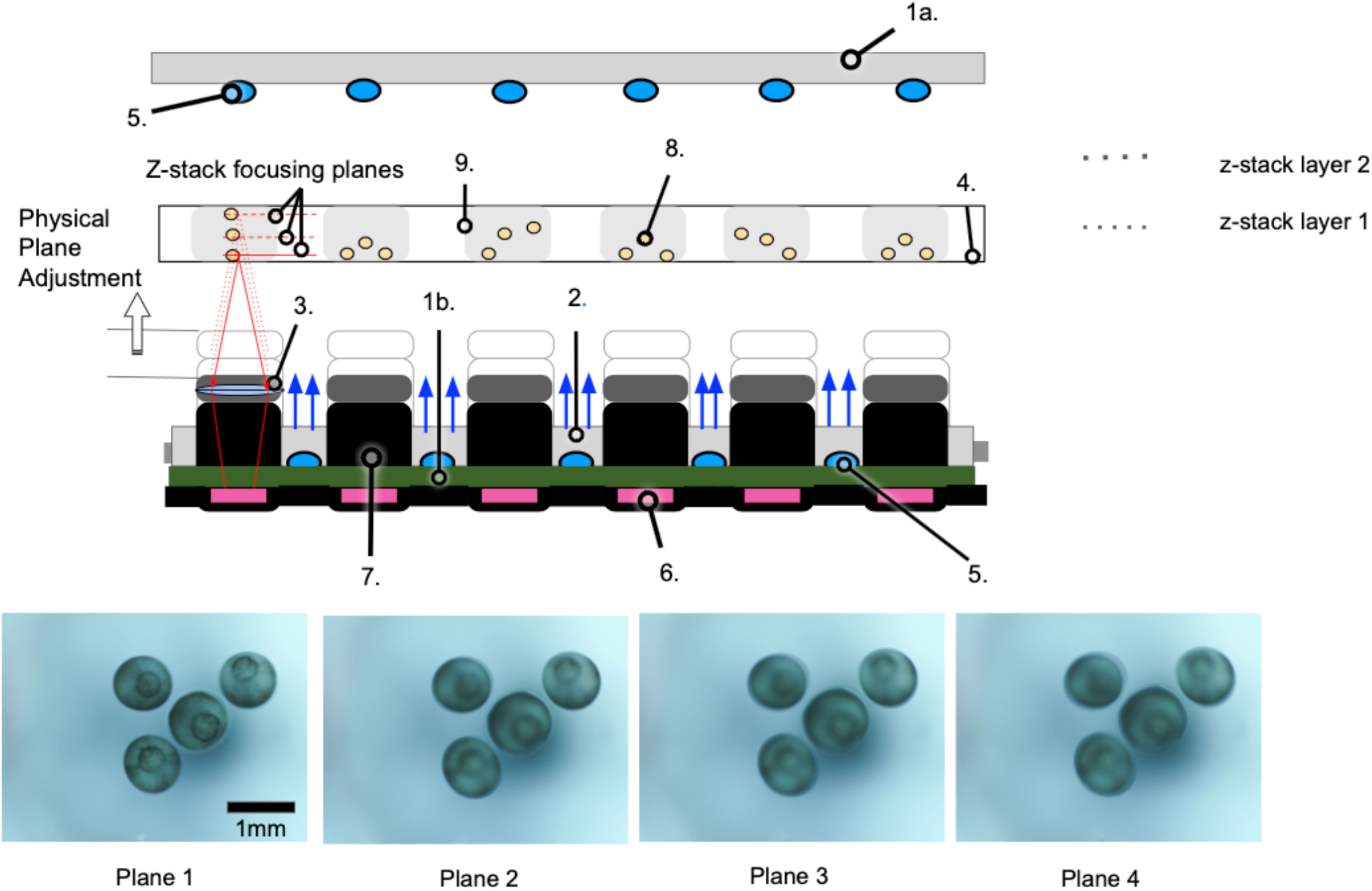
Schematic representation of the z-stack function. 1.a = Over-the-plate illumination board, 1.b = Under-the-plate illumination board, 2 = Acrylic Light Diffuser, 3 = Lenses, 4 = Cell Culture Plate, 5 = LEDs, 6 = Raspberry Spy Cameras, 7 = 3D Printed Camera Bodies, 8 = Biological Sample (e.g., Frog Embryos), 9 = Individual Culture Well. The photos at the bottom were taken at four planes 0.3 mm apart. The blastopore is only in focus in Plane 1

**Figure 4.**
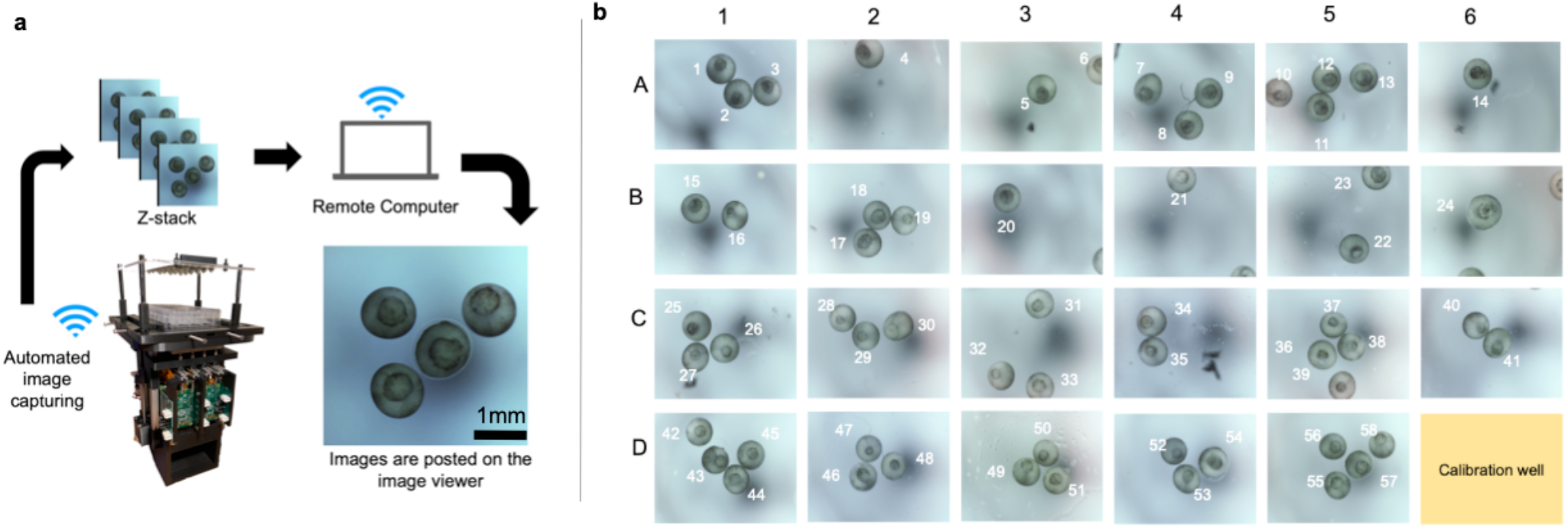
System architecture: **A** The images are autonomously collected and wirelessly transferred to a remote computer for viewing or post processing **B** Image of 23 wells observing 57 frog embryos

### Longitudinal imaging of *Xenopus tropicalis* embryonic development

As proof of principle of the longitudinal live imaging capabilities of the Picroscope, we imaged the development of *Xenopus tropicalis* embryos from the onset of gastrulation through organogenesis (Figure 4B, 5 and 6). The fertilization and development of *Xenopus* occurs entirely externally, which allows scientists to easily observe and manipulate the process^41^. For decades, *Xenopus* have been heavily used in biology studies to model a variety of developmental processes and early onset of diseases, particularly those of the nervous system^42^. While several species of *Xenopus* are used in different laboratories around the world, *Xenopus tropicalis* is one of the preferred species due to its diploid genomic composition and fast sexual maturation^43, 44^. Normal development and optimal husbandry of *Xenopus tropicalis* occurs at 25-27°C,^45, 46^closely approximating standard room temperature, which eliminates the need of special environmental control for most experiments.

**Figure 5.**
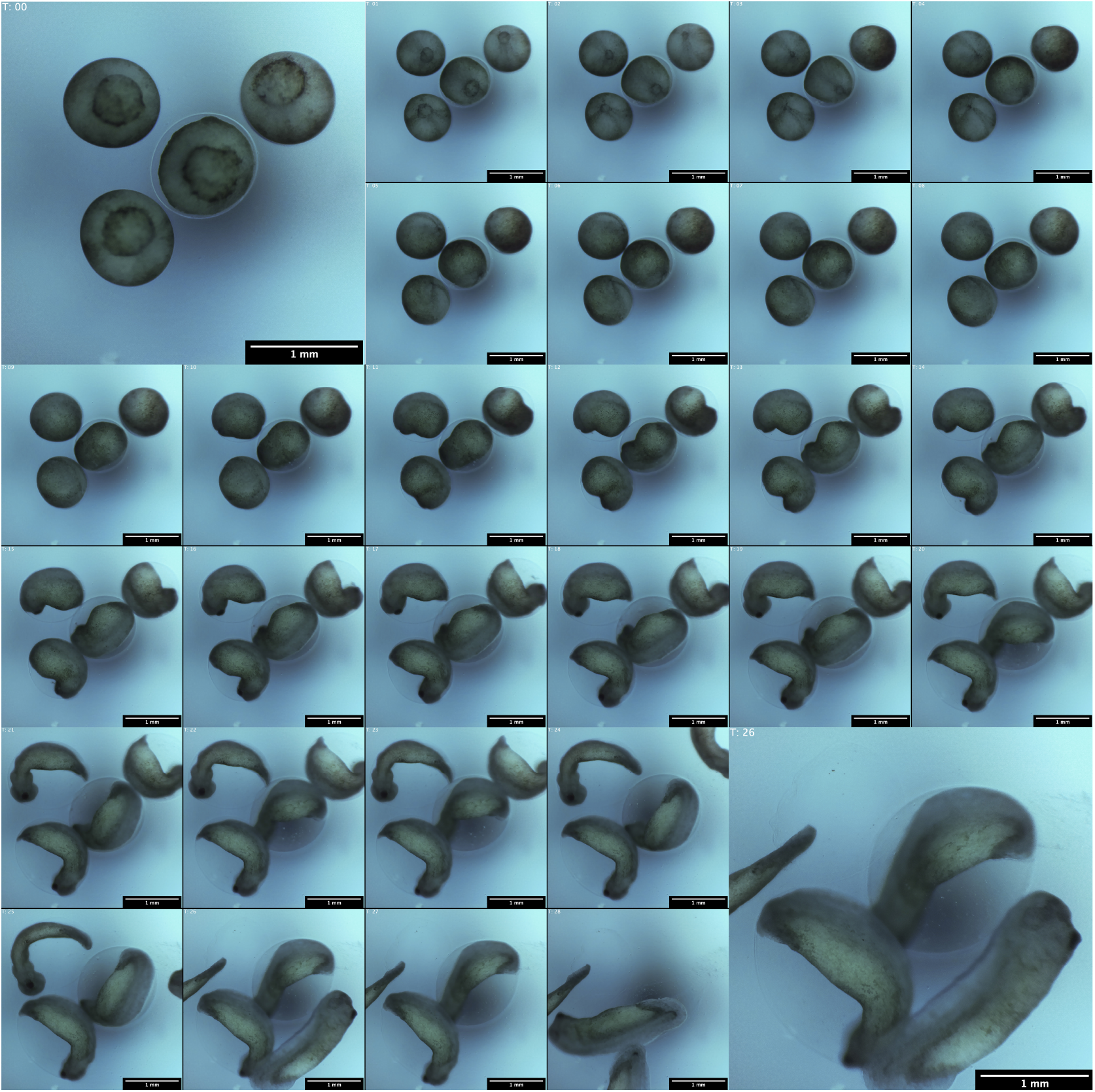
Longitudinal imaging of Xenopus tropicalis development. Images of a representative well in which 4 frog embryos developed over a 28 hours period. Images were taken hourly. White Balance adjusted for visibility

**Figure 6.**
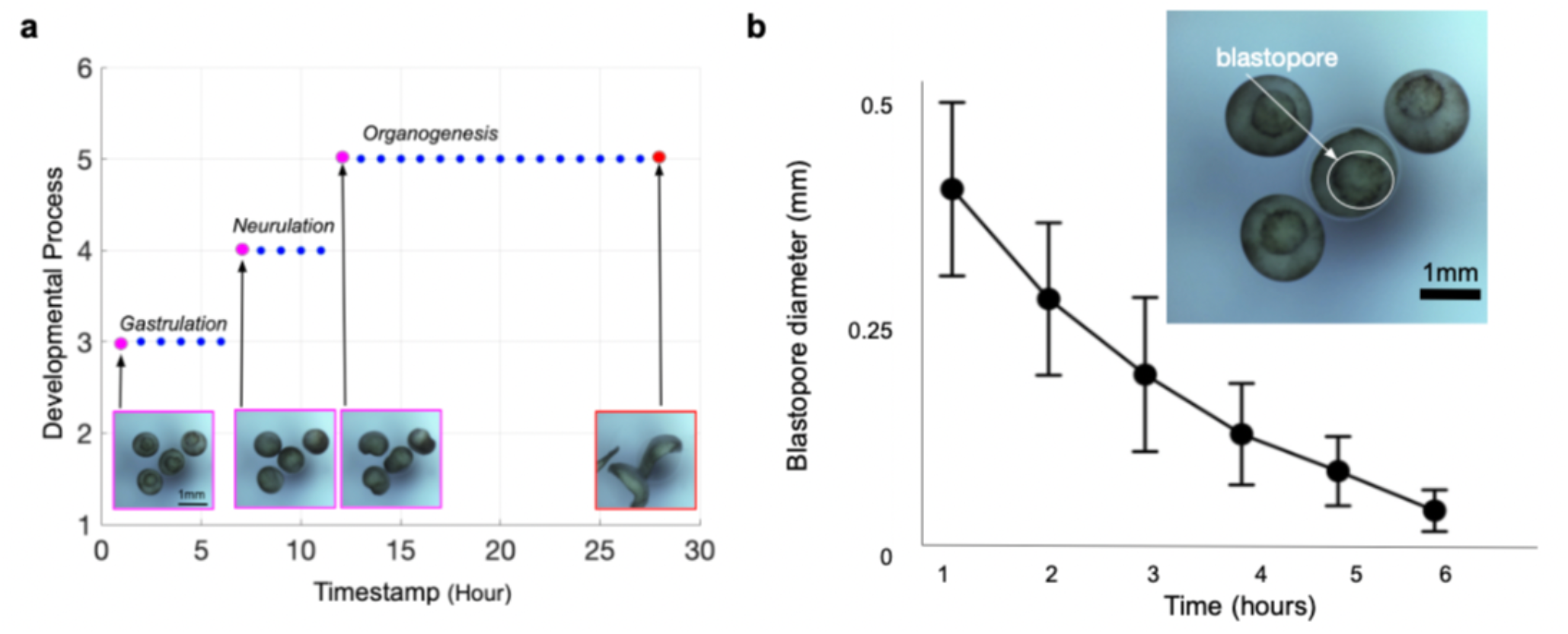
Longitudinal imaging allows the tracking of individual developmental processes: **a** The images shown in figure 5 were taken hourly over a 28 hours period and encompass 3 developmental stages: Gastrulation, neurulation and organogenesis. Y-Axis represents the stages of frog embryonic development: 1 = Fertilization, 2 = Cleavage, 3 = Gastrulation, 4 = Neurulation, 5 = Organogenesis, 6 = Metamorphosis. X-Axis represents the timepoint at which it occurs. Each dot in the plot represents a timepoint in which the images were taken. Magenta = the beginning of each developmental process. Red = the end of the experiment at 28 hours. Blue = intermediate timepoints. **b** Diameter of the blastopore is reduced over time from gastrulation to neurulation. Top right panel shows an example of an individual blastopore. A total of 27 embryos were considered for the analysis.

Given these convenient experimental advantages and their large size, *Xenopus* embryos have been used extensively to understand the development of the vertebrate body plan, with particular success in elaborating the complex cellular rearrangements that occur during gastrulation and neural tube closure^47, 48^. These experiments rely on longitudinal imaging of developing embryos, often at single-embryo scale with dyes, fluorescent molecules, and computational tracking of single cells^48, 49^. These studies have elucidated key cellular mechanical properties and interactions critical to vertebrate development, often replayed and co-opted during tumorigenesis. There exists an opportunity to scale these experiments to be more high-throughput with the Picroscope, as one could image hundreds of developing embryos simultaneously, rather than having to move the objective from embryo to embryo during development, or repeating the experiment many times.

We imaged *Xenopus tropicalis* embryos over a 28 hour time period. Four embryos were placed in each of the 23 wells used in a 24-well plate, and we used an extra well as calibration (Figure 4b and 5). The embryos were grown in simple saline solution and the experiment took place at room temperature. Imaging was performed hourly starting at gastrulation (Figure 6). Then, we visually inspected each image and mapped the embryos to the standard stages of frog development, categorizing their development in gastrulation, neurulation and organogenesis (Figure 6a). Finally, we took a subset of 27 embryos and measured the diameter of the blastopore as the embryos underwent gastrulation (Figure 6b). We observed a progressive reduction of blastopore diameter over a 6 hour time period, consistent with progression through gastrulation and the start of neurulation. This simple experiment demonstrated that the Picroscope can be used for longitudinal sequential imaging and tracking of biological systems.

### In-incubator imaging of human embryonic stem cells and brain organoids

While many biological systems including zebrafish, planaria and frogs develop at room temperature and atmospheric gas concentrations, mammalian models require special conditions requiring an incubator enclosure. Mammalian models include 2D monolayer cell cultures, as well as 3D organoid models of development and organogenesis. They have been used to assess molecular features and effects of drugs for a variety of phenotypes including cell proliferation,^50, 51^ morphology^52, 53^and activity^54, 55^, among others.

Deploying electronics and 3D printed materials inside tissue culture incubators presents some unique challenges. The temperature and humidity conditions can cause electronics to fail and cause certain plastics to offgas toxins^56^. Plastics can also be prone to deformation in these conditions. A common solution for protecting electronics and preventing offgassing is to use inert protective coatings e.g., Parylene C. This requires expensive clean room equipment. Instead we print all of the components with Polylactic acid (PLA), a non-toxic and biodegradable material, to prevent deformation we print using 100% infill and reinforce vulnerable elements with aluminum MakerBeam profiles. We coat all electronic components with Corona Super Dope Coating to protect the electronics from the conditions (heat and humidity) of an incubator.

We tested the functionality of the Picroscope inside a standard tissue culture incubator by imaging 2D-monolayers of human embryonic stem cells (hESCs) (Figures 7a-b). To demonstrate the capacity of our system to perform longitudinal imaging across the z-axis, we imaged human cerebral cortex organoids embedded in Matrigel (Figure 7c). Using this system we could monitor the growth of the organoids, as well as the outgrowth of neuronal processes (Figure 7d). Tracking of individual cells within organoid outgrowths allowed us observe their migration patterns and behavior (Figure 7d). Altogether, we show the feasibility of using our system for longitudinal imaging of mammalian cell and organoid models.

**Figure 7.**
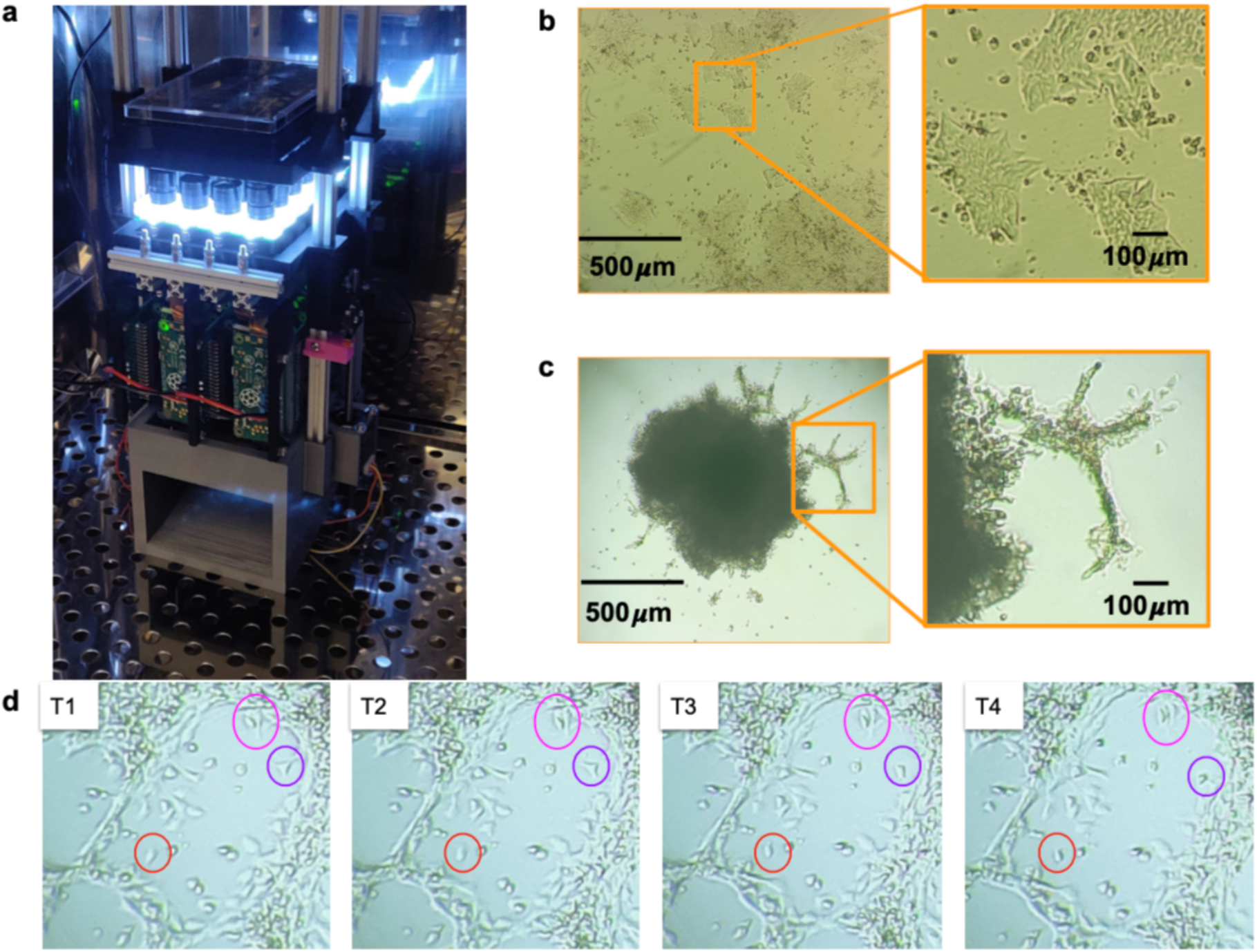
In-incubator imaging of mammalian cell and cortical organoid models. **a** The Picroscope inside a standard tissue culture incubator. **b** Imaging of human embryonic stem cells as a model of 2D-monolayer cell cultures. **c** Longitudinal imaging of human cortical organoids embedded in Matrigel. Zoomed images show the outgrowth of neuronal processes originating in the organoids. **d** Manual Longitudinal tracking of migration of individual cells in embedded cortical organoids over 40 minutes. Images were taken every 10 minutes.

## DISCUSSION

The combination of 3D printed technology and open-source software has significantly increased the accessibility of academic and teaching laboratories to biomedical equipment^57^. Thermocyclers, for example, were once an expensive commodity unattainable for many laboratories around the world^58^. Now, low cost thermocyclers have been shown to perform as well as high-end commercially available equipment^59^. Inexpensive thermocyclers can be used in a variety of previously unimaginable contexts, including conservation studies in the Amazon^60^, diagnostics of Ebola, Zika and SARS-CoV-2^60, 61^, teaching high school students in the developing world^10^and epigenetic studies onboard the International Space Station^62^.

Simultaneous imaging of biological systems is crucial for drug discovery, genetic screening, and high-throughput phenotyping of biological processes and disease^50, 51, 53^. This technique typically requires expensive multicamera and robotic equipment, making it inaccessible to most. While the need for a low cost solution has long been appreciated^63^, few solutions have been proposed. Currently, the low cost solutions can be grouped in two categories: 1) those that use of gantry systems that move an individual camera through multiple wells, performing “semi-simultaneous” imaging^33–35^ or 2) those that use acquisition of large fields of view encompassing multiple wells (usually with limited resolution per well, followed by post-processing of images^24, 64^. Neither of these solutions is optimal to perform true simultaneous imaging of biological replicates across multiple conditions. To overcome these limitations, the Picroscope performs an automated image capture of a standard 24 wells cell culture plate using 24 individual objectives. The images are then transferred to a remote computer or server (using the Picroscope’s internet connection), where they can be viewed and/or processed (Figure 4), with minimal intervention.

Commercial electronic systems for simultaneous imaging of biological samples are typically designed to image cells plated in monolayers^65^. Yet, significant attention has been given to longitudinal imaging-based screens using whole organisms. These have included zebrafish^66, 67^, worms^68^, and plants^69^. Many times, the results of the screens are based on single plane images or in maximal projections obtained from external microscopes^68, 69^. The Picroscope was designed to overcome these limitations and image along the z-axis. This is accomplished with fine adjustment by two stepper motors that lift the elevator unit that holds all 24 camera objectives (Figure 3).

To date, few 3D printed microscopes are designed to function inside incubators^24, 36^. We have run the picroscope in the incubator for three weeks. This makes the Picroscope compatible with screens in 3D mammalian models including organoids^70, 71^. We have shown a proof of principle of this function by performing longitudinal imaging of human cortical organoids and analyzing the behavior and movement of individual cells (Figure 7).

We anticipate many useful applications of the Picroscope and derivatives of it. Here we demonstrated the versatility of the Picroscope across animal and cell models in different environmental conditions. The modular nature of the system, allows for new features to be easily built and added. For example, defined spectrum LED light sources and filters for fluorescent imaging would enable longitudinal studies of the appearance and fate of defined sub populations of cells in a complex culture by taking advantage of genetically encoded fluorescent reporter proteins^66^. Similarly, the use of fluorescent reporters or dyes that respond to dynamic cell states such as calcium sensors allow long-term imaging of cell activity^54^. The Picroscope, paves the way towards increased accessibility and democratization of multi-well multi-week simultaneous imaging experiments in diverse biological systems.

## MATERIALS AND METHODS

### 0.1 System Setup

The picroscope uses 1 Raspberry pi 4 as a hub pi that is connected to an ASUS N300 Router via ethernet. The 24 Raspberry Pi 0Ws communicate with via wifi in order to transmit the photos being taken by each of the 24 Zero spy cameras. Each Raspberry pi device, requires a micro sd card where a copy of our code can be flashed using Balena Etcher. For the lenses on the Zero Spy Cameras, we use Arducam 1/2” M12 Mount 16mm Focal Length Camera Lenses. An Arduino Uno with the Adafruit V2 Motor Driver Shield is used to control both Nema 11 External 34mm Stack 0.75A Lead 0.635mm/0.025” Length 100mm motors. To detect the bottom of the preset z-stack, we use a limit switch mounted on the 3D printed elevator component via two M2.5 fasteners. All of the constructive elements of the picroscope were designed using Fusion360 or AutoCAD computer aided design (CAD) packages. The 3D printed compoents are sliced with the Prusaslicer with 100% infill, and with 0.15mm quality preset settings. The 3D printed components were manufactured using a Prusa MK3S 3D printer. The printing material is black PLA. MakerBeam aluminum extrusion elements were used (1) as a structural component for the elevator piece and (2) as a guide for the vertical sliding stages. The sliding stages were constructed using 4 200mm 10×10mm MakerBeam Aluminum extrusions. The custom electronics were designed on a standard 1.6 mm FR4 two-layer PCB. The Power distribution board PCB (Figure 2 c 11) is designed to power and provide structural support for the Raspberry Pi Zero W through their 5V GPIO pins. This design is modular and allows us to have a double-sided PCBs and the same design can be used for the two single-sided PCBs. The relay board (Figure 2 e) allows us to trigger the illumnation boards individually as well as shut off power to the entire picroscope in the event of temperature overheat condition.

For bright field microscopy, the over head light PCB (Figure 2 a 1) uses MEIHUA white LEDs with a brightness of 228 450MCD, and the brightness can be adjusted through a potentiometer. The PCB for lighting from below (Figure 2a 4), are NCD063W3 Chip LEDs. All custom PCBs are manufactured by PCBWay (China); the cost, including shipping is approximately $ 2 per board. All electronic components were purchased from Digi-Key Electronics (MN, United States). The PCBs were populated and assembled at the University of California at Santa Cruz shown in Figure 2. All electronic components (Raspberry Pi, Arduino, PCBs) were coated with Corona Super Dope Coating to shield the hardware from the effects of condensation due to elevated humidity inside incubator environments. We used a Nomad883 pro to CNC a custom diffuser made out of frosted acrylic.

### 0.2 Biological Samples

#### Animals

All animal experiments complied with the regulations of the University of California, Santa Cruz and the University of California, San Francisco.

#### Frogs

*Xenopus tropicalis* husbandry was performed as previously described^72^. Adult animals were maintained and cared for according to established IACUC protocols. Animals were wild type and both sexes were used. Animals were ovulated using human chorionic gonadotropin (Sigma-Aldrich, C1063) according to Sive et al. (2000)^73^ and both *in vitro* fertilizations and natural matings were used. Embryos were maintained in 1/9 modified Ringer’s solution^73^ and staged according to Nieuwkoop and Faber (1958)^74^. Blastopore size was measured in ImageJ/FIJI (NIH) and plotted in GraphPad Prism software version 9.

#### Zebrafish

Fertilized zebrafish eggs were purchased from Carolina Biological Supply Company (Catalog # 155591) and maintained in media containing 15 mM sodium chloride (Sigma-Aldrich, S9888), 0.5 mM potassium chloride (Sigma-Aldrich, P3911), 1 mM calcium chloride dihydrate (Sigma-Aldrich, 223506), 1 mM magnesium sulfate heptahydrate (Sigma-Aldrich, 1058822500), 150 µM potassium phosphate monobasic (Sigma-Aldrich, P5655), 50 µM sodium phosphate dibasic heptahydrate (Sigma-Aldrich, S9390), 0.7 mM sodium bicarbonate (Sigma-Aldrich, S5761) and 0.1% methylene blue (Sigma-Aldrich, M9140).

#### Planaria

*Dugesia tigrina* Brown planaria worms were purchased from Carolina Biological Supply Company (Catalog # 132954). Planaria were grown in bottled water (Poland Spring). Water was changed every other day.

#### Human embryonic stem cells and cortical organoids

All hESC experiments used the H9 cell line (WiCell)^75^. hESCs were grown on vitronectin (Thermo Fisher Scientific, A14700) coated plates and cultured using StemFlex Medium (Thermo Fisher Scientific, A3349401). Passages were performed incubating the cells in 0.5mM EDTA (Thermo Fisher Scientific, 15575020), in DPBS for 5 minutes.)

To generate cortical organoids, we first dissociated hESCs into single cells and re-aggregated them in Aggrewell 800 well plates (STEMcell Technologies) at a density of 3,000,000 cells per well with 2mL of Aggrewell Medium (STEMcell Technologies) supplemented with Rho Kinase Inhibitor (Y-27632, 10 µM, Tocris, 1254) (Day 0). The following day (Day 1), we supplemented the aggregates with WNT inhibitor (IWR1-ε, 3 µM, Cayman Chemical, 13659, Days 1-10) and TGF-βinhibitor (SB431542, Tocris, 1614, 5 µM, days 0-10). On Day 2, aggregates were transferred by pipetting out of the Aggrewell plate with wide bore P1000 pipette tips onto a 37 µm filter and then transferred to ultra low adhesion 6-well plates. Media was changed on Days 4, 8 and 10, by replacing 2mL of conditioned media with fresh media. On Day 11 the medium was changed to Neuronal Differentiation Medium containing Eagle Medium: Nutrient Mixture F-12 with GlutaMAX supplement (DMEM/F12, Thermo Fisher Scientific, 10565018), 1X N-2 Supplement (Thermo Fisher Scientific, 17502048), 1X Chemically Defined Lipid Concentrate (Thermo Fisher Scientific, 11905031) and 100 U/mL Penicillin/Streptomycin supplemented with 0.1% recombinant human Fetal Growth Factor b (Alamone F-170) and 0.1% recombinant human Epidermal Growth Factor (R&D systems 236-EG). On Day 12, the organoids were transferred in 90 µL media to a custom glass-PDMS microfluidic chip for imaging/feeding containing 50 µL Matrigel hESC Qualif Matrix (BD 354277) bringing the total volume in the well to 120 µL. Partially embedding the organoid in Matrigel in this way led to 2D outgrowths on the surface of the Matrigel. Feeding occurred automatically every hour replacing 30 µL Neuronal Differentiation Medium.

## SUPPLEMENTAL MATERIALS

**Supplemental Video S1: Video Imaging of a Developing Zebrafish**

## ACKNOWLEDGMENTS

This work is supported by the Schmidt Futures Foundation SF 857. Research reported in this publication was also supported by the National Institute Of Mental Health of the National Institutes of Health under Award Number R01MH120295 and the National Science Foundation under award number NSF 2034037. We would like to thank Jeremy Linsley and Wiktoria Leks for providing us zebrafish for this study. In addition, we would like to thank Arnar Breevoort for providing experimental support. H.R.W. was supported by grant U01MH115747-01 from NIMH to Matthew State. M.A.M.-R. was partially supported by grant TL1 TR001871 from the NIH National Center for Advancing Translational Sciences. K.V. was supported by grant T32HG008345 from the National Human Genome Research Institute (NHGRI), part of National Institutes of Health (NIH), USA. S.R.S was supported by NIMH award. M.T was support by NSF EAGER. D.H was supported by Schmidt Futures Foundation.

## AUTHOR CONTRIBUTIONS

V.T.L., P.V.B., P.P., K.V. worked on hardware, software, and assembly of the Picroscope. Y.R., E.A.J. worked on early prototypes. V.T.L., P.V.B, H.R.W., G.L.M., S.S., J.M.R., A.A.P., M.A.M.-R. performed Biological experiments. H.R.W., A.A.P., T.J.N., S.R.S., M.A.M.-R., D.H., M.T. conceived the experiments.D.H., M.T., S.R.S, A.A.P. M.A.M.-R., M.R. and T.J.N. supervised the team and secured funding. V.T.L., P.V.B., E.A.J, M.A.M.-R., M.T. wrote the manuscript with contributions from all authors.

## COMPETING INTERESTS

The authors have written patents covering the technology described in this article. AAP is in the board of Herophilus. DH is a Howard Hughes Medical Institute Investigator. The authors declare no other conflict of interest.

## Notes

### Competing Interest Statement

The authors have written patents covering the technology described in this article. AAP is in the board of Herophilus. The authors declare no other conflict of interest

